# Tribe: The collaborative platform for reproducible web-based analysis of gene sets

**DOI:** 10.1101/055913

**Authors:** René A. Zelaya, Aaron K. Wong, Alex T. Frase, Marylyn D. Ritchie, Casey S. Greene

## Abstract

**Background:** The adoption of new bioinformatics webservers provides biological researchers with new analytical opportunities but also raises workflow challenges. These challenges include sharing collections of genes with collaborators, translating gene identifiers to the most appropriate nomenclature for each server, tracking these collections across multiple analysis tools and webservers, and maintaining effective records of the genes used in each analysis.

**Description:** In this paper, we present the Tribe webserver (available at https://tribe.greenelab.com), which addresses these challenges in order to make multi-server workflows seamless and reproducible. This allows users to create analysis pipelines that use their own sets of genes in combinations of specialized data mining webservers and tools while seamlessly maintaining gene set version control. Tribe’s web interface facilitates collaborative editing: users can share with collaborators, who can then view, download, and edit these collections. Tribe’s fully-featured API allows users to interact with Tribe programmatically if desired. Tribe implements the OAuth 2.0 standard as well as gene identifier mapping, which facilitates its integration into existing servers. Access to Tribe’s resources is facilitated by an easy-to-install Python application called *tribe-client*. We provide Tribe and tribe-client under a permissive open-source license to encourage others to download the source code and set up a local instance or to extend its capabilities.

**Conclusions:** The Tribe webserver addresses challenges that have made reproducible multi-webserver workflows difficult to implement until now. It is open source, has a user-friendly web interface, and provides a means for researchers to perform reproducible gene set based analyses seamlessly across webservers and command line tools.

## Background

Modern bioinformatics webservers provide users with ready access to powerful analytical approaches, but users that attempt to integrate analyses across one or more servers encounter reproducibility challenges. Workflows comprised of many steps and different servers and command-line tools run the risk of losing track of what genes were used in each step.

We designed Tribe to address three major challenges that plague reproducible analysis with multi-webserver pipelines: cross-server sharing, identifier mapping, and version control. Tribe is open source and can be installed within a company, bioinformatics core, or other facility. We also provide a public instance that contains gene information for 9 organisms with 16 different types of gene identifiers.

Specifically, when running analyses in different webservers using a set of genes thought to be involved in a biological process or pathway, there is danger of missing genes from large lists of genes that are copied and pasted onto the web interfaces of these webservers. PILGRM [1], GeneWeaver [2], GeneMania [3], and GeNets [4], for example, are webservers built to perform analyses on user-specified genes or gene sets. PILGRM and GeneMania find new genes similar to a given input set of genes using various data (in the case of PILGRM, a second set of genes saved as a negative standard is also used). GeneWeaver and GeNets allow users to analyze sets of genes using different tools through their web interfaces. However, none of these webservers provide an API that can be accessed either programmatically by individual researchers or by other webservers that run analyses. Therefore, researchers analyzing gene sets on multiple webservers must copy and paste their collections of genes and hope that webservers support common formats and identifiers.

This raises an additional challenge: webserver developers may choose to implement different identifiers for genes - translating between these identifiers presents an additional, error-prone step. Some webservers or analysis tools for analysis of human genes may use traditional HGNC [5] gene symbols as identifiers to run an analysis, while others may use Entrez [6] or Ensembl [7]. In some cases, specific genome locations map to more than one gene. In this case, whether one gene or both genes are used as the identifier for the location is inconsistent between different tools/webservers. If researchers are sharing their collections of genes with collaborators through e-mail or scattered communications methods, the risk for identifier translation errors is further increased. The multitude of gene identifiers and difficulty tracking versions hinders reproducible science, as the set of genes used in one step of the workflow may unintentionally differ from those used in another step.

The Tribe webserver was built to address these challenges: it allows researchers to input collections of genes and to use them across multiple servers. A “collection” is a group of genes that are connected (optionally through publications) to user-defined processes, pathways, or analytical results (Figure 1). Researchers can build their collections, create new versions of them, and share them with their collaborators. Tribe keeps track of every time a new version of a collection is created, and every time a collection is “forked”, or copied, to be further versioned. Tribe’s database maps between gene identifiers to provide collections to webservers using the gene identifiers that the webserver requests. Tribe has an easy-to-use web interface for users without programming expertise. This interface is implemented as a JavaScript application that accesses a fully featured API. For programmatically inclined users, this API provides direct access to all functionality available to the web interface.

**Figure 1.**
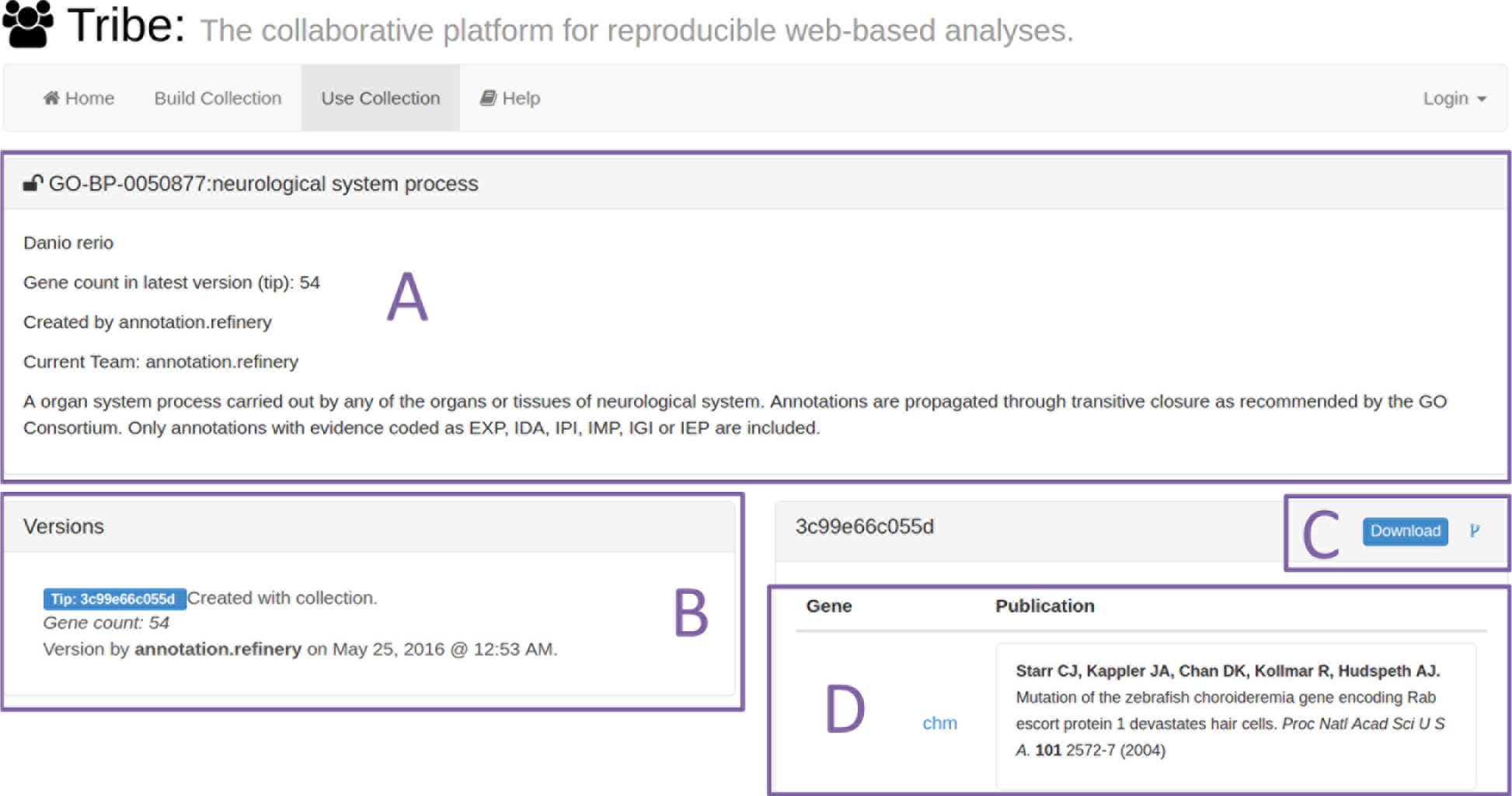
Example screenshot of the Tribe webserver. (A) Collection metadata is provided, which includes the organism, creator, and a description. Example public collections are populated with experimentally-supported annotations from the Gene Ontology [32]. (B) Versions are tracked and shown in this panel. The tip version, used by some webservers, is marked in blue in the interface. (C) Users can download this version of the collection or fork it to create their own copy that they can build upon through Tribe. (D) Genes associated with the collection and, optionally, publications that led to their annotation in this collection are shown in this panel.

## Implementation

Tribe is an open-source webserver, which uses a relational database to store information about organisms, genes, users, and webservers. Here we highlight the general properties of the webserver. We also provide a publicly available instance of Tribe, which is described in more detail in “*Our Publicly Available, Free Tribe Instance*”.

Back-end server functionality is implemented using the Django web framework [8] and written in Python (currently running and unit tested on Python v2.7.12). The web interface is built using a combination of JavaScript via the AngularJS framework [9], HTML, and CSS. Database access is handled through standard Django queries allowing Tribe to be deployed to any relational database system supported by Django. Tribe implements full text search of genes and gene sets using Haystack [10], a modular search package for Django, so a Haystack-supported search backend is also required. We provide Fabric [11] functions that can install Tribe’s requirements on an Amazon Web Services Ubuntu image, and another set within the Tribe repository to deploy an instance of Tribe to a machine that has all required software packages installed.

While Tribe’s database contains a number of database tables, the following are the most relevant to Tribe’s gene collection and versioning functionality:

- Genes – Tribe implements loading of cross-reference identifiers from files provided by the Entrez database. These include but are not limited to: Entrez [6], Ensembl [7], HGNC [5], HPRD [12], MGI [13], MIM [14], SGD [15], UniProtKB [16], TAIR [17], WormBase [18], RGD [19], FLYBASE [20], ZFIN [21], Vega [22], IMGT/GENE-DB [23], and miRBase [24]. Tribe uses Entrez to define a gene, and Tribe can map genes between any of these different types of gene identifiers. Tribe uses symbols for display purposes, and it provides user- or server-requested identifiers for external analyses.
- Publications – Tribe saves publication objects by saving its citation information (Pubmed [25] ID, title, authors, date, journal, volume, pages, and issue). Users can add publications to genes in their collections as evidence for the genes being part of the collection. Users are able to add new publication objects to the database by simply entering a Pubmed ID into the user interface.
- Collections – Tribe saves version-controlled collections of genes with, optionally, publications used to associate a gene to a collection. Users can update collections, and Tribe will preserve a history of all of the versions saved by a user (Figure 2). Collection versioning is implemented by generating a 40-character SHA-1 hash of the genes, publications, and parent of each version. This implementation mirrors the technique for version control used by Mercurial [26]. The 40-character SHA-1 hash provides a version identifier that external webservers can use to provide access to results using the specific version of a gene list or to save a new version after analyses. For webservers that implement version-aware analyses, users can run an analysis using any previously saved version of their gene collection. Collections can be shared with collaborators, who can create new versions of the collection if granted permission by the collection creator (Figure 3).

**Figure 2.**
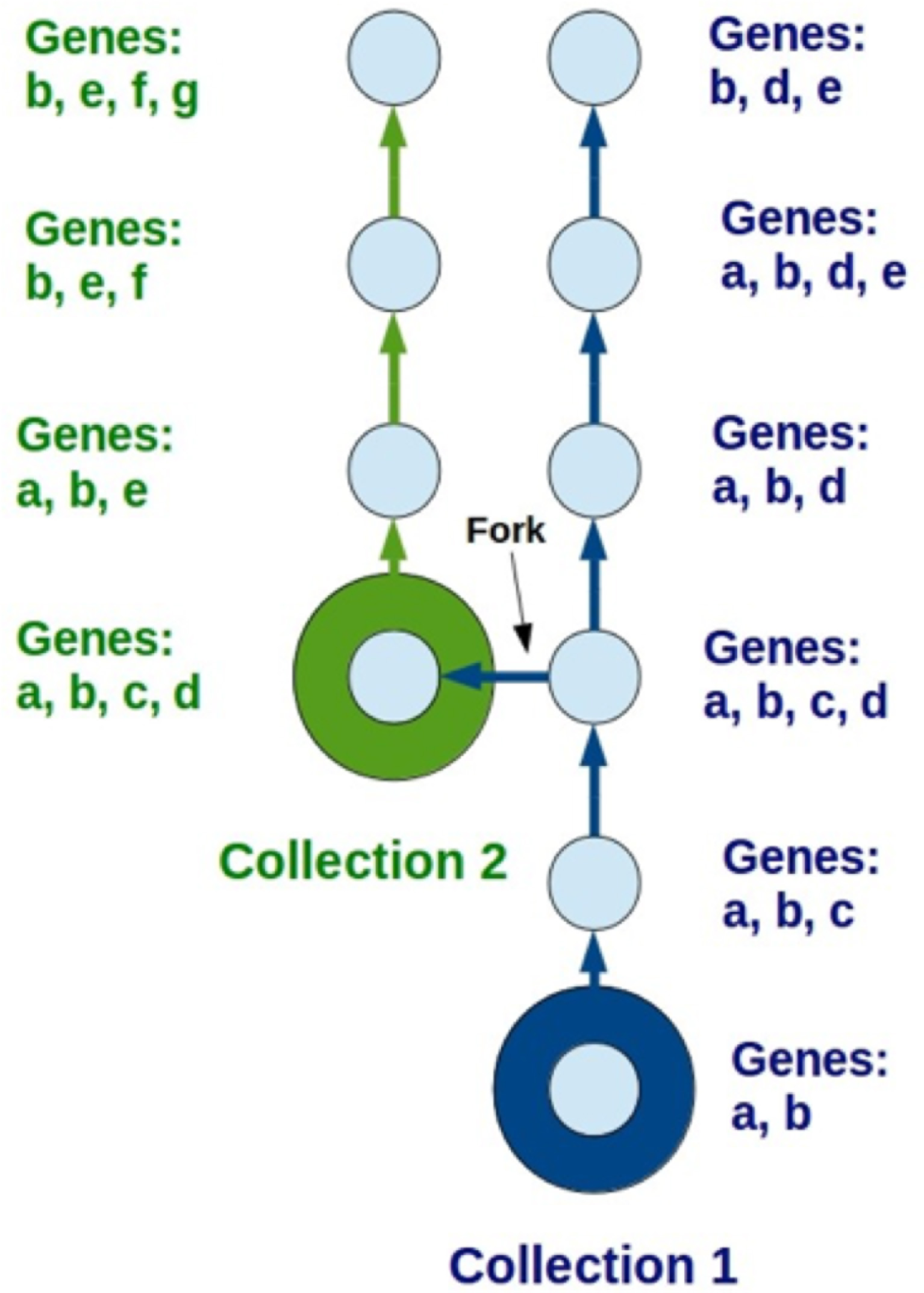
Tribe supports versioning and forking of collections. The blue and green circles represent where a new collection is being created. Tribe preserves the history of all of the collection’s versions and their contents. Furthermore, collections can be copied into new collections, or “forked”, at any point and versioned thereafter. This can be done either by the original collection’s author or an approved collaborator.

Aside from genes, publications, and collections, the Tribe database contains other tables that enable the webserver’s functionality. Some tables, such as the user sessions, are automatically created by Django [8] to provide common webserver functionality, whereas others are specific to Tribe functionality. For example, there is a table for the different organisms supported, and genes in the database have a foreign key linking them to their corresponding organism. Finally, there are tables that keep track of the different collaborations between users and their collections in Tribe.

Tribe is accessible both programmatically and also through a web interface. Tribe has a REST API that serves its resources (collections, versions, genes and publications) as JSON objects, which can be consumed by client servers, applications and even command-line tools. Currently, Tribe is integrated into the GIANT [27], ASD [28], FNTM [29] and WISP (in preparation) webservers (Figure 4). LOKI [30, 31], which parses sets of genes annotated to pathways and processes (such as GO terms [32] and KEGG pathways [33]), produces JSON output and can be configured to send this output to Tribe’s API using the OAuth 2.0 protocol [34] (Figure 4). This allows users to access these sources of curated pathways via both the web interface and a standardized programmatic interface.

### Tribe-client

We provide a self-contained Python package to facilitate the access to Tribe’s resources. Tribe-client is installable in a user’s Python environment, where they can make use of tribe-client’s functions to programmatically get and create Tribe resources. These functions include all the code necessary to make the HTTP requests to the Tribe API endpoints so that the users only need to specify the parameters of the request. If a webserver developer wants to connect their webserver to Tribe, they can do so by installing tribe-client on their webserver, and make use of the functions that automatically handle authentication via the OAuth 2.0 protocol [34]. Furthermore, if the webserver is built using the Django framework [8], the integration with tribe-client will be seamless, as tribe-client also provides views and templates for these webservers.

## Discussion

### Version-controlled Collections

Researchers who use Tribe will typically be either bench biologists or bioinformaticians who are accustomed to performing analyses across multiple webservers or software packages. They will often have a base of knowledge that can be summarized as one or more sets of genes. For example, these could be the genes that lead to a phenotype of interest during a screen, genes that are members of a particular pathway of interest, genes identified during a literature search, or genes that result from an analytical workflow.

Tribe helps these users maintain a version-controlled history of each set of genes, regardless of where they originate. These gene sets, termed “collections” in Tribe, can be used directly with integrated webservers or software. Users grant selected webservers or software access to their collections. These webservers or software packages can read directly from Tribe and can write results back to the server. This allows users and integrated webservers to record the input and output versions associated with such gene set based analyses. Throughout the process, users and approved collaborators can add genes to their collections and may optionally also add publications that support why a certain gene is part of the set of genes (Figure 3). Tribe saves a versioned history of both genes and publications, and these versions can be used directly through its API. If desired, users can download a tab-delimited file containing all the genes and publications in a specific version with a selected type of gene identifier. Our current version of Tribe is designed to simplify the usage and management of gene sets, though the system may be extended to other types of collections.

**Figure 3.**
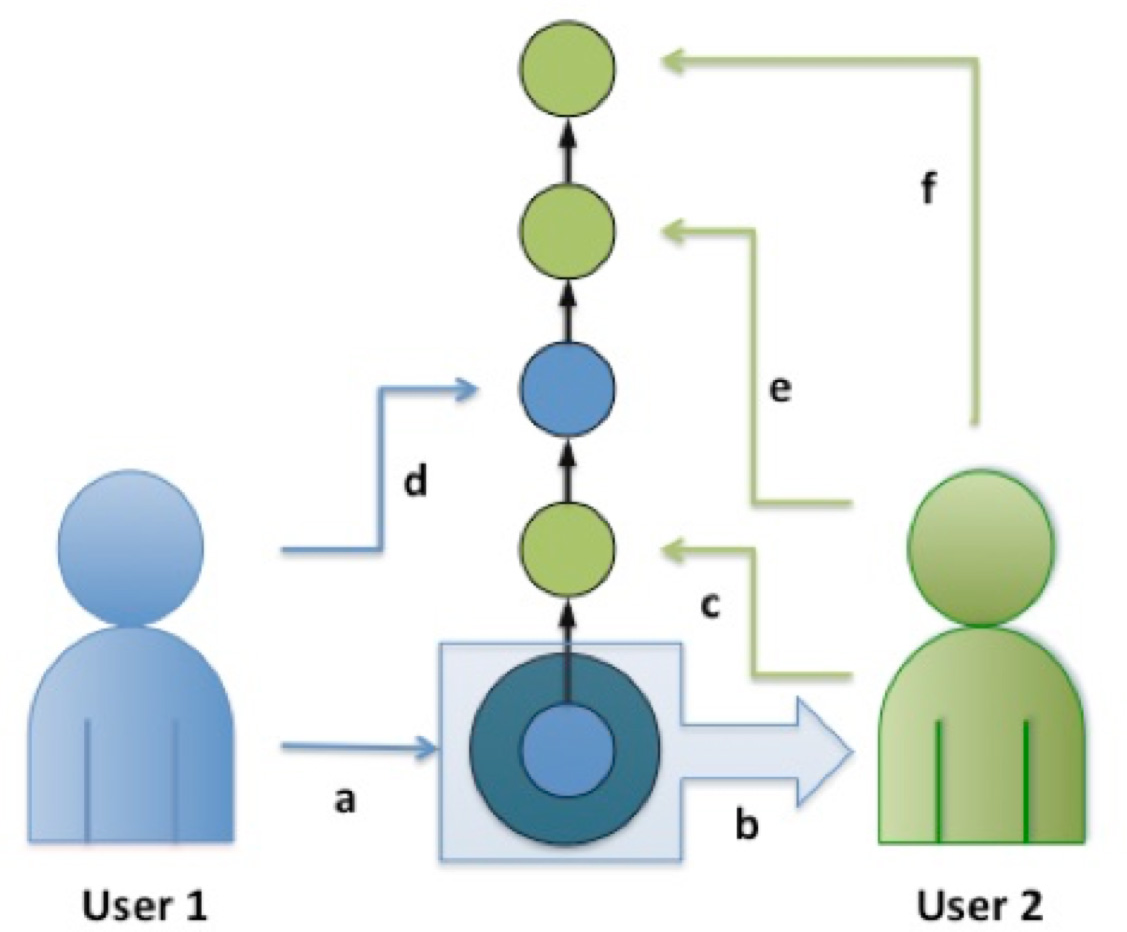
Tribe allows users to share their collections. In this example, User 1 a) creates a collection with certain genes and publications, and b) shares it with User 2. User 2 then c) adds and/or removes members in the collection based on literature curation or experimental results to create a new version. Afterwards, User 1 and 2 create a few more versions of the collection based on their findings as the research project proceeds (d, e, f).

### Fully Featured Temporary User Accounts

Users can view publicly available Tribe collections, whether programmatically or through the web interface. As with many servers, users can create an account to store and share collections and versions by entering an email address and selecting a password.

Tribe also provides a feature, called temporary anonymous accounts, for users who want to create one or more collections for analysis but who do not wish to enter an e-mail or password. A temporary anonymous account allows users to create, edit and delete their own collections, and these users can perform analyses on integrated webservers. Users are connected to their temporary account via a cookie, so they may only access these resources from the computer and Internet browser they used to create the anonymous user account. Temporary anonymous account holders can access their account and resources for 1 year, until they erase their browser cookies, or they log out of the account in that browser - whichever comes first. At any point before the account expires, these users can convert a temporary anonymous account into a full, regular Tribe account by entering an email address and a password. All resources saved up to that point are maintained in the new, regular account, where they can access and modify them.

This feature also allows webservers that integrate with Tribe to be fully usable with no registration requirement, even if they need to take user-generated collections as input or to save user-specific output.

### Collaborative and Reproducible Workflows

Full user accounts allow users to share collections and invite collaborators via email addresses. When running analyses in client webservers connected to Tribe using tribe-client, e.g. GIANT [27], users can allow GIANT to access their collections via Tribe’s OAuth 2.0 [34] authentication framework (Figure 4). This provides access to the Tribe collections that they have access to, i.e. their own and those that they collaborate on, within the external webserver. External webservers can implement read, write, or read-write and the user is notified which permissions are requested. This allows webservers to implement use cases where user collections are read and analytical results are written back either as new versions or new collections.

**Figure 4.**
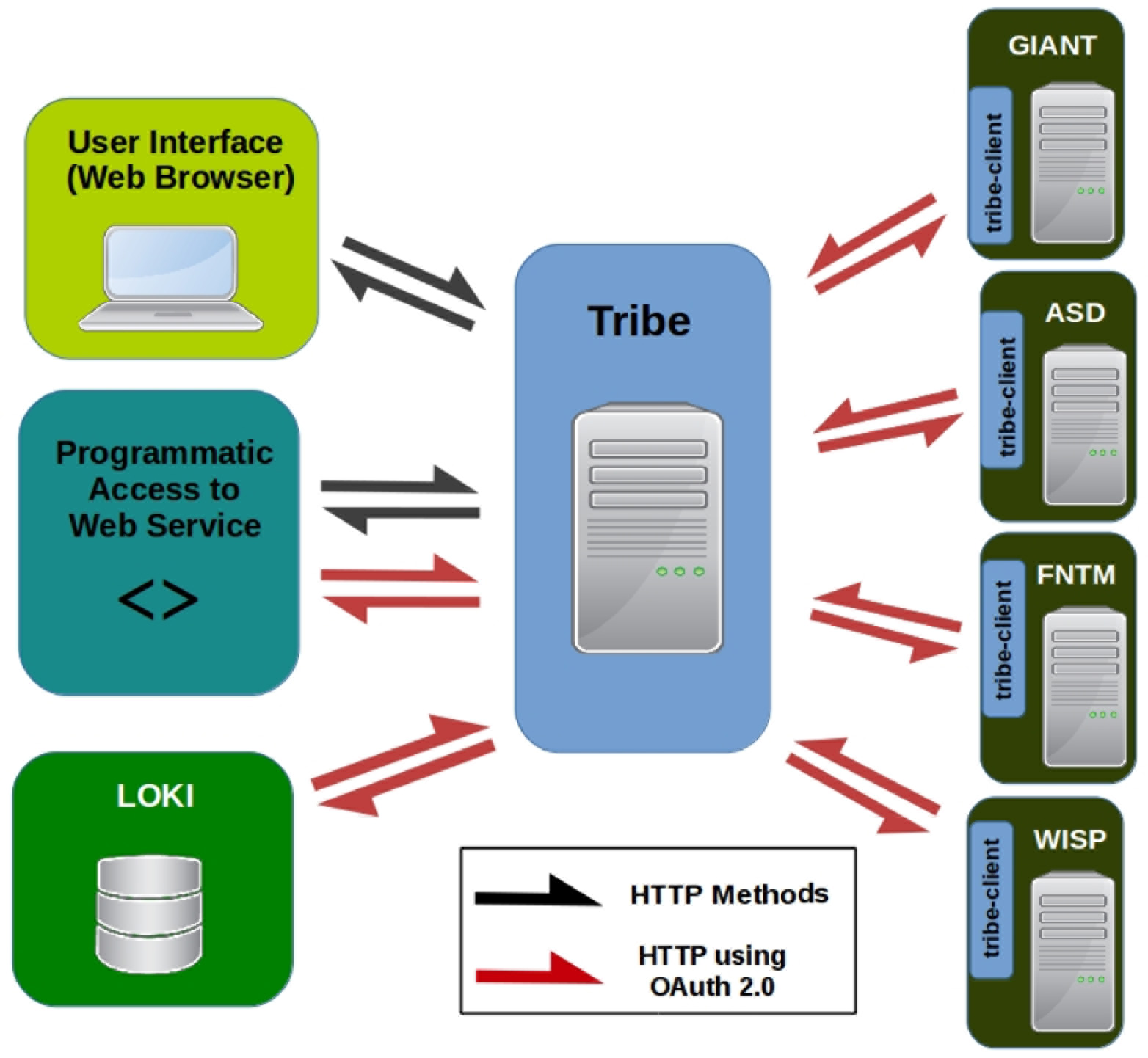
Tribe access diagram. Users can access their resources by navigating Tribe’s web interface at the main Tribe website, but they can also access Tribe by writing scripts that make HTTP requests to Tribe’s API. If users are using the web interface, they can login directly in the Tribe website to access their private resources, or they can authenticate using the OAuth 2.0 protocol [34] if they wish to access their private resources programmatically. Collections can be made public, and no authentication is required for access to public collections. LOKI [30, 31], GIANT [27], ASD [28], FNTM [29] and WISP (in preparation) currently connect to Tribe via the OAuth 2.0 protocol. We also provide “tribe-client”, an open-source application (used by LOKI, GIANT, ASD, FNTM and WISP) to facilitate the connection of outside webservers and analytical tools to Tribe resources.

Tribe separates the functionalities of running an analysis from storing the collections used as input or output. This modularity facilitates reproducible user-created workflows that combine different webservers or software packages. Instead of writing software to specifically integrate each potential subsequent analysis, server developers only need to integrate with Tribe via a standard OAuth 2.0 mechanism. In this way, a complex analysis pipeline that involves several different steps and is prone to errors can be simplified by using OAuth capabilities provided by Tribe as a common framework.

### Our Publicly Available, Free Tribe Instance

The publicly accessible instance of Tribe is running on an Amazon Web Services Ubuntu image and is available at https://tribe.greenelab.com. This instance uses the relational database system PostgreSQL [35] and Elasticsearch [36] as the backend search engine for Haystack. It currently contains the gene information for 9 organisms in its database: *Homo sapiens*, *Mus musculus*, *Drosophila melanogaster*, *Rattus norvegicus*, *Caenorhabditis elegans*, *Saccharomyces cerevisiae*, *Danio rerio*, *Arabidopsis thaliana* and *Pseudomonas aeruginosa*. It also has the 16 different types of gene identifiers previously mentioned loaded in its database to accommodate the needs of many different webservers and analysis tools. We are adding organisms and identifiers as users and webserver developers request them.

### Multi-server Analysis Example Case Study

The following case study illustrates the advantages of using Tribe for a multi-server workflow. In this case study, a researcher is investigating genes involved in some way with Autism Spectrum Disorder (ASD) with her collaborator.

Firstly, she queries the set of genes that is part of Copy-Number Variant (CNV) 7q11.23 in the ASD webserver [28], and copies the list of genes from the web interface onto a text file or spreadsheet and saved as the genes that are part of that CNV. Afterwards, her collaborator wants to analyze this set of genes in the context of a tissue network from the GIANT webserver [27], so she sends him the file with the list of genes. The collaborator copies and pastes the list of genes onto the GIANT web interface to run an analysis using the global tissue network. He copies and pastes the output of this analysis onto another text or spreadsheet file, which he sends back to the first investigator.

Using Tribe this process is simplified and recorded. The first researcher queries the ASD webserver for a CNV and clicks the “Save query genes to Tribe” button in the query result page (Figure 5A). The set of genes is sent to Tribe, and she may enter a description for the collection if desired. She can share this collection with her collaborator by clicking the “Add collaborator to team” button (Figure 5B). The collaborator can log into GIANT using Tribe and this collection will be selectable in the “Gene Set” drop-down menu on GIANT’s home page (Figure 5C). By clicking search, he’ll reach the analysis results page (Figure 5D), which provides the option to save the results of the network query back to Tribe. These results can then be shared with the first investigator. Tribe and its integrated webservers handle any necessary identifier mapping, and the process of webserver-to-Tribe interactions requires the input and output of each step to be recorded.

**Figure 5.**
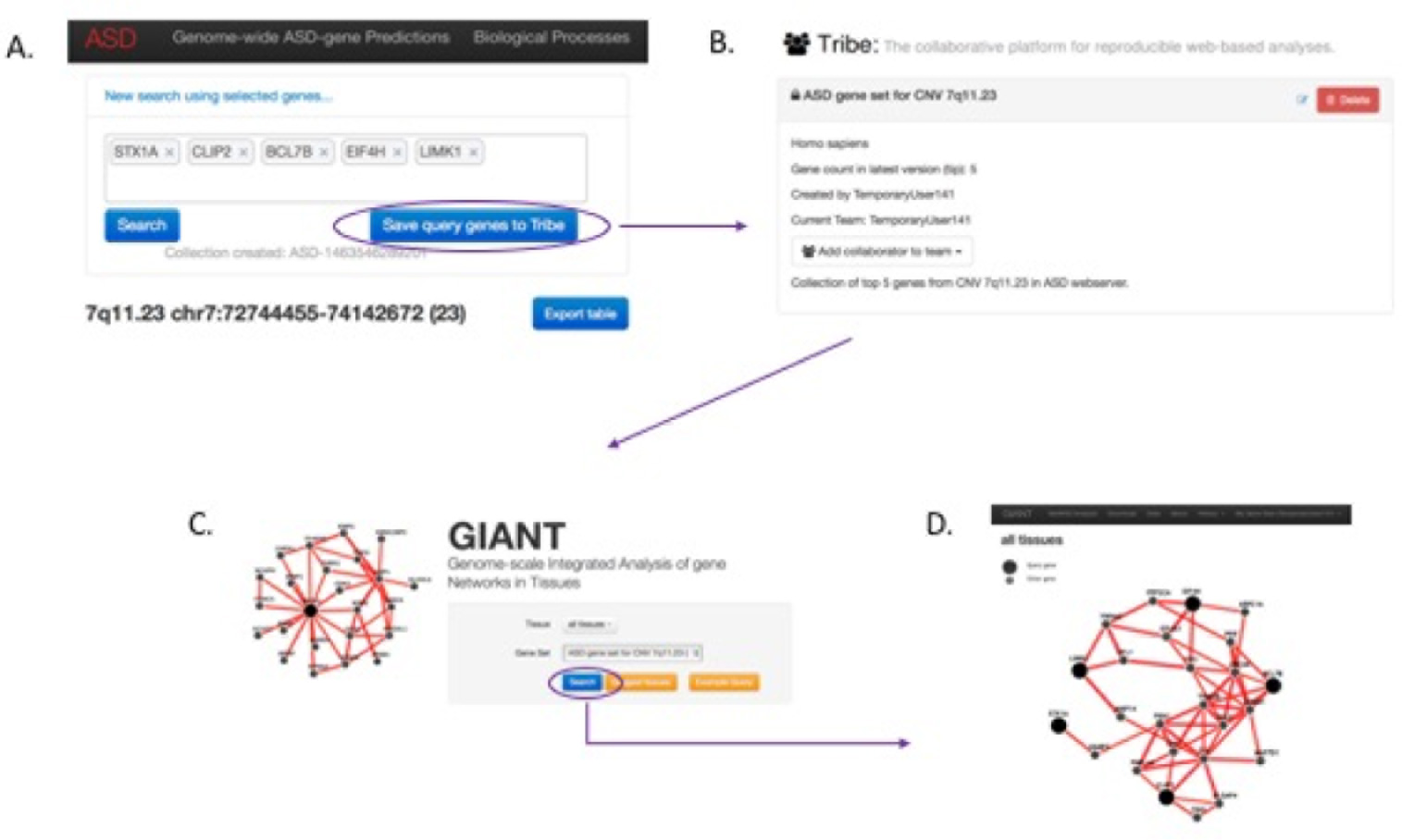
Case study workflow. The user’s path through the servers is shown with purple arrows. A researcher queries genes that are part of the desired CNV in the ASD webserver [28] and saves them to Tribe as a new collection (A, B). She then performs an analysis using this set of genes with the global tissue network in the GIANT webserver [27] (C, D).

## Conclusions

Tribe is a flexible, multifaceted webserver that is accessible by other webservers, analytical software, and its web interface. As webservers and other analysis tools become specialized for certain types of analyses, Tribe helps developers link their webservers with other servers that provide complementary analytical capabilities. This helps researchers link these tools together to create comprehensive analysis pipelines with capabilities beyond those of any single server or tool.

Tribe not only helps servers and tools communicate with each other, it also helps researchers collaborate with each other by allowing them to curate and manage collections of genes together. Sharing gene lists, especially when they are very large, is cumbersome via e-mail, and all of the genes in the list must be translated into identifiers that analysis tools can interpret. Furthermore, Tribe keeps a clear record of a collection’s version history so that users, or other researchers, can reproduce their analysis.

For bioinformatics developers, Tribe provides user authentication capabilities via OAuth 2.0 [34], eliminating the need for account management software to be implemented in each server, greatly simplifying development. Tribe is an open source project. Its development is ongoing and we are continuing to add support for additional organisms, identifiers, and client capabilities. Tribe fits into a growing ecosystem of servers that provide fast APIs for computational biologists. For example, the recently published server MyGene.info uses a fast API to provide comprehensive information about individual genes [37]. For users, Tribe provides a means to collaborate on reproducible workflows across multiple servers and software packages.

## Declarations

### Ethics approval and consent to participate

Not applicable

### Consent for publication

Not applicable

### Availability of data and materials

Our instance of Tribe can be accessed at https://tribe.greenelab.com using any web browser supported by AngularJS [9] or via its API. There is no login requirement to access any public resources. Users can create a temporary anonymous account, which does not require an email address or password, to create resources in lieu of a full account. These temporary accounts can be converted to a full account at any time. The Tribe webserver, source code to facilitate deployment, and tribe-client software package are available under the permissive 3-clause BSD license. The source code for Tribe and tribe-client can be found at https://github.com/greenelab/tribe and https://github.com/greenelab/tribe-client, respectively. Tribe-client can be installed via the Python Package Index (PyPI).

## Authors’ contributions

RAZ and CSG designed the webserver and client application. RAZ and CSG developed the webserver and client application. RAZ, AKW and ATF extensively tested the webserver and client application. RAZ drafted the manuscript. AKW, ATF, MDR and CSG critically revised the manuscript.

## Acknowledgements

Not applicable

